# Transcriptomic changes during the replicative senescence of human articular chondrocytes

**DOI:** 10.1101/2023.11.07.565835

**Authors:** Aysegul Atasoy-Zeybek, Gresin P. Hawse, Christopher V. Nagelli, Consuelo Lopez De Padilla, Matthew P. Abdel, Christopher H. Evans

## Abstract

Osteoarthritis (OA) is a degenerative joint disease and a leading cause of disability worldwide. Aging is a major risk factor for OA, but the specific mechanisms underlying this connection remain unclear. Although chondrocytes rarely divide in adult articular cartilage, they undergo replicative senescence *in vitro* which provides an opportunity to study changes related to aging under controlled laboratory conditions. In this pilot study, we performed bulk RNA sequencing on early- and late-passage human articular chondrocytes to identify transcriptomic changes associated with cellular aging. Chondrocytes were isolated from the articular cartilage of three donors, two with OA (age 70-80 years) and one with healthy cartilage (age 26 years). Chondrocytes were serially passaged until replicative senescence and RNA extracted from early- and late-passage cells. Principal component analysis of all genes showed clear separation between early- and late-passage chondrocytes, indicating substantial age-related differences in gene expression. Differentially expressed genes (DEGs) analysis confirmed distinct transcriptomic profiles between early- and late-passage chondrocytes. Hierarchical clustering revealed contrasting expression patterns between the two isolates from osteoarthritic samples and the healthy sample. Focused analysis of DEGs on transcripts associated with turnover of the extra-cellular matrix and the senescence-associated secretory phenotype (SASP) showed consistent downregulation of Col2A1 and ACAN, and upregulation of MMP19, ADAMTS4, and ADAMTS8 in late passage chondrocytes across all samples. SASP components including IL-1α, IL-1β, IL-6, IL-7, p16^INK4A^ (CDKN2A) and CCL2 demonstrated significant upregulation in late passage chondrocytes originally isolated from OA samples. Pathway analysis between sexes with OA revealed shared pathways such as extracellular matrix (ECM) organization, collagen formation, skeletal and muscle development, and nervous system development. Sex-specific differences were observed, with males showing distinctions in ECM organization, regulation of the cell cycle process as well as neuron differentiation. In contrast, females exhibited unique variations in the regulation of the cell cycle process, DNA metabolic process, and the PID-PLK1 pathway.

## Introduction

Osteoarthritis (OA) is the most common form of arthritis and a leading cause of disability among older people. Risk factors for OA include aging, obesity, genetic predisposition, hormonal changes, anatomical abnormalities, and prior joint injury.^1^ Of these, aging is the most prominent contributor to OA risk.^2,3^ The prevalence of OA rises with age, with about 22% of people over 40 and one in three over 65 affected. Women are disproportionately impacted^4,5^ reflecting the exponential increase in the prevalence of OA after menopause (≥ 45 years).^6^

Replicative senescence refers to the irreversible cell cycle arrest that occurs after a finite number of cell divisions *in vitro*. This loss of replicative capacity was first described by Hayflick and Moorhead,^7,8^ who found that primary human fibroblasts in culture underwent a progressive loss of mitotic capacity with serial passage, leading to irreversible cell cycle arrest after approximately 30-40 population doublings. The inability of primary cells to proliferate beyond a finite number of doublings is now known as the “Hayflick limit”. Evans and Georgescu^9^ first demonstrated that cultures of articular chondrocytes from rabbits, dogs, and humans underwent replicative senescence and suggested that chondrocyte senescence explains the association of idiopathic OA with age. Because chondrocytes in adult cartilage rarely divide, even in OA, it was suggested that aging *in situ* reflected extended periods in the G_0_ phase of the cell cycle.^10^

Multiple recent studies of articular cartilage have documented age-related declines in chondrocyte cell yields, reduced ability to proliferate, with enhanced degradation of components of the extracellular matrix (ECM) particularly aggrecan, the major proteoglycan of cartilage.^11–13^ Such changes contribute to cartilage thinning, mechanical weakening and ultimately loss of cartilage function.^14^ Chondrocytes derived from older individuals exhibit characteristic features of cellular senescence, including shortened telomeres, heightened cell mortality, elevated senescence-associated β-Galactosidase (SA-β-Gal) activity, and increased expression of key regulators such as p53, p21^CIP1^, and p16^INK4A^(CDKN2A).^15–18^ Moreover, they display an excessive production of reactive oxygen species (ROS).^19^ Elevated production of ROS triggers the activation of genes associated with chondrocyte dedifferentiation and senescence.^20,21^

Senescent chondrocytes adopt a senescence-associated secretory phenotype (SASP).^22,23^ This phenotype involves the secretion of a range of factors, including pro-inflammatory cytokines^24,25^ like IL-1, IL-6, IL-7, IL-8, IL-17, OSM, GM-CSF, and TNFα, as well as matrix metalloproteinases (MMPs)^26,27^ such as MMP1, MMP3, MMP10, and MMP13, MMP19, and other proteinases that degrade the ECM of cartilage, including ADAMTS4, ADAMTS5, and ADAMTS8 alongside altered activity and expression of growth factors^28–32^ including TGF-β and IGF-1. Notably, there is evidence of elevation in SASP factor levels with age in OA patients.^3,12,33^ These age-related alterations in cartilage physiology are likely contributors to the onset and progression of OA.^34^

In this pilot study, we harvested chondrocytes from the knees of three individuals and compared the transcriptomes of early and late passage cells. Two of the human chondrocyte cultures were established from older individuals (72 and 80 years old) with OA, and one from a young healthy adult (26 years old). Chondrocytes from all donors were serially passaged to replicative senescence^9^ and bulk RNA-sequencing was preformed to identify age-related transcriptomic changes.

## Materials and Methods

### Study Design

As summarized in figure 1, chondrocytes recovered from knee articular cartilage were serially passaged until they reached their Hayflick limit. RNA was extracted from early passage (passage 1) and late (terminal) passage cultures and subjected to bulk sequencing and bioinformatic analysis. The abundance of certain transcripts was confirmed by quantitative RT-PCR, and in the case of IL-1, protein expression was confirmed by ELISA. The quality of cartilage formed in pellet culture was assessed at passages P (1) and P (18).

**Figure 1.**
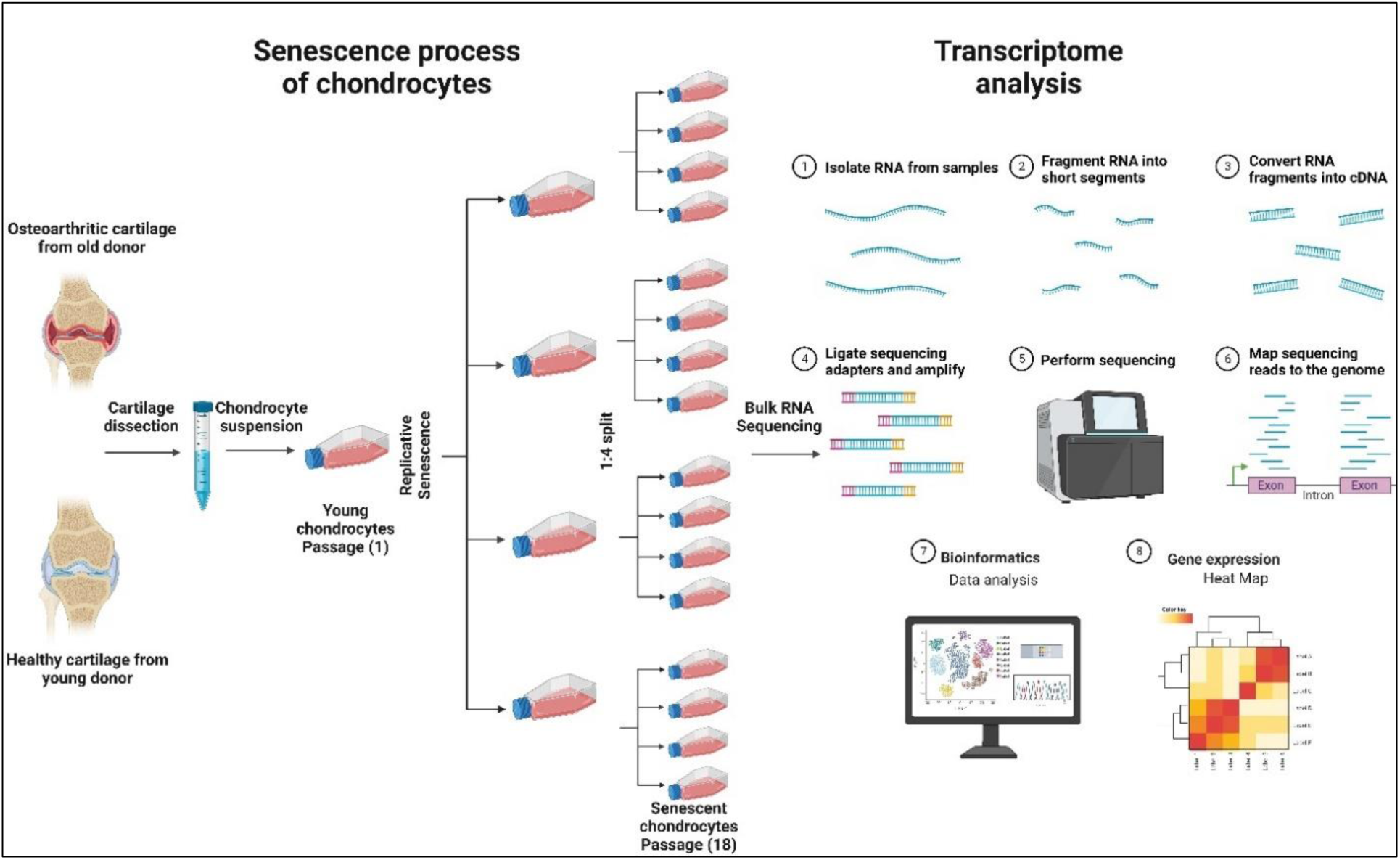
Overview of the study including replicative senescence of the chondrocytes and bulk RNA sequencing steps.

### Biospecimen Processing

#### Cartilage collection

Human osteoarthritic cartilage biospecimens (n=2, patient 1: male aged 80, patient 2: female aged 72) were collected from individuals undergoing total knee arthroplasty at Mayo Clinic, Department of Orthopedic Surgery with approval by the Institutional Review Board (IRB number: 13-005619).

#### Chondrocyte Isolation

Healthy human articular chondrocytes were purchased from Lonza (Basel, Switzerland; Catalog number: CC-2550), (n=1, male aged 26). For the two samples recovered from patients undergoing total knee arthroplasty, full-thickness pieces of articular cartilage were removed aseptically from the femoral condyles and tibial plateaux using a scalpel. The cartilage was cut into 1-2 mm^3^ pieces and washed with phosphate buffered saline (PBS), (Gibco, Gaithersburg, MD, USA). The cartilage pieces were digested for 2 hours at 37°C in 0.2 % pronase (Millipore Sigma, St. Louis, MO, USA; Catalog number: 10165921001) in Dulbecco’s Modified Eagle Medium/Nutrient Mixture F-12 (DMEM/F-12; Sigma-Aldrich, St. Louis, MO, USA) with a mixture of penicillin (100 U/ml) and streptomycin (100 μg/ml) (Gibco) (1% P/S). After pronase digestion, the chondrocytes were isolated by further digestion for 16 hours at 37°C in 0.05 % type II collagenase (Gibco, catalog number: 17101015) in DMEM/F-12 media supplemented with 5% fetal bovine serum (Gibco) (FBS) and 1% P/S. Digestion was performed in a Labnet Problot™ 6 hybridization oven (Labnet Int Inc., Cary, NC, USA). Subsequently, the digestion solution was filtered through a 70 μm cell strainer and cells were washed. Cells were counted, seeded into 75 cm^2^ culture flasks at a density of 1.0 x 10^4^ cells/cm^2^ in 10 mL of DMEM/F12 supplemented with 1% P/S and 10% FBS, and incubated at 37° in 5% CO_2_.

### Serial passage of chondrocytes

Cells were recovered from confluent cultures by trypsinization and re-seeded into 75 cm^2^ flasks at a 1:4 split ratio. Briefly, the cells were treated with 4 mL of TrypLE™ (Thermo Fisher Scientific, Waltham, MA, USA) and incubated at 37° in 5% CO_2_ for 5 minutes to detach the cells. Subsequently, 8 mL of medium containing 10% FBS was added, followed by centrifugation at 400 g for 5 minutes. The supernatant was then discarded, and the pellet was reconstituted in 1 mL of medium. Cell counting was performed, and 250 μL of medium was seeded into a 75 cm² flask. This process was continued until cells could no longer achieve confluence, at which point they were considered to have reached their Hayflick limit. This occurred after 36 doublings in all three cultures.

### 3D Pellet culture

Early or late (terminal) passage chondrocytes were cultured as pellets as previously described.^35^ Briefly, the cells were plated at a density of 2 × 10^5^ cells per well in polypropylene, v-bottom, 96- well plates (Corning, Corning, NY, USA). The plates were centrifuged at 400 g for 5 minutes, supernatants were removed and replaced with chondrogenic medium (high glucose DMEM (Sigma-Aldrich), 1% insulin-transferrin-selenium premix (BD Biosciences, Franklin Lakes, NJ, USA), 100 nM dexamethasone (Sigma-Aldrich), 50 µg/mL ascorbate-2-phosphate (Sigma-Aldrich), 1 mM sodium pyruvate (Sigma-Aldrich), 40 µg/mL L-proline (Sigma-Aldrich), and 10 ng/mL TGFβ1 (PeproTech, Cranbury, NJ, USA; Catalog no: 100-21). During the initial 24 hours of incubation, cells formed free-floating aggregates. Media were changed the following morning and every other day thereafter.

### Histology

#### Pellet cultures

Pellets were fixed for 1 hour with 4% paraformaldehyde, centrifuged at 400 rpm for 5 minutes, then washed with 95% alcohol. The cell pellets were resuspended in 100 µL of 0.8% low melting point agarose solution (Invitrogen Carlsbad, CA, USA; Catalog No: 16520-050), followed by re-centrifugation on a table-top centrifuge at 1500 rpm for 5 minutes. The resulting compact agarose cell pellet button was allowed to solidify for 30 minutes at 4 °C. The solidified compact agarose cell was placed in a biopsy bag folded into a cassette and processed and embedded as per routine histology. 5-µm-thick sections were cut using an automatic microtome (HM 355S, Thermo Fischer Scientific) and mounted onto positively charged slides (Superfrost™ Plus Microscope Slides, Thermo Fisher Scientific).

#### Toluidine blue staining

Briefly, 5-µm-thick paraffin-embedded sections were deparaffinized and rehydrated through a series of xylenes and graded alcohols to distilled water, followed by staining with 0.1% toluidine blue pH 2.3 for 1.5 minutes, rinsed in distilled water for 1 minute, then quickly dehydrated in graded ethyl alcohol series, cleared with xylene, and mounted with xylene-based mounting medium (Richard-Allan Scientific™, Kalamazoo, MI, USA). Bright field images were acquired using an automated inverted microscope (Olympus IX83) at 4X and 20X magnifications using the cellSens Olympus imaging software.

### Bulk RNA sequencing and data analysis

#### RNA isolation

A two-step RNA isolation protocol was followed for total RNA extraction: Organic phase isolation was performed with TriReagent®/ bromo–3–chloropropane (BCP) (Thermo Fisher Scientific/Molecular Research Center, Inc., Cincinnati, OH, USA) and then used for total RNA isolation using a RNeasy Plus Mini Kit (Qiagen, Germantown, MD, USA) according to the manufacturer’s instructions. RNA concentration was measured using a nanophotometer® (Implen, Inc., Westlake Village, CA, USA). RNA purity was A_260/280_ and A_260/230_ ˃ 2.0. RNA integrity was assessed using an Agilent 2100 bioanalyzer (Agilent, Santa Clara, CA, USA) (RIN ˃ 8).

#### Illumina Stranded mRNA library prep with Illlumina NextSeq2000 Sequencing

RNA libraries were prepared using 200 ng of total RNA in 20 µl according to the manufacturer’s instructions for the Illumina Stranded mRNA Ligation Sample Prep Kit (Illumina, San Diego, CA). The concentration and size distribution of the completed libraries were determined using an Agilent Bioanalyzer DNA 1000 chip and Qubit fluorometry (Invitrogen). Libraries were sequenced using Illumina’s standard protocol for the NovaSeq 6000. The NovaSeq SP PE100 flow cell was sequenced as 100 X 2 paired end reads using NovaSeq Control Software v1.8.0. Base-calling was performed using Illumina’s RTA version 3.4.4.

#### Bioinformatics and data analysis

Fastq files for all samples were processed using the Mayo RNA-Seq bioinformatics pipeline, MAP-RSeq version 3.1.4.^36^ MAP-RSeq employs a fast, accurate and splice-aware aligner, STAR^37^ for generating bam files by aligning sequencing reads to the reference human genome build hg38. Gene and exon expression quantification was performed using the Subread^38^ package to obtain both raw gene counts and normalized Fragments Per Kilobase per Million mapped reads (FPKM) values. Finally, FASTQC^39^ and MultiQC^40^ tools were used for comprehensive quality control assessment of the aligned reads. Results from all modules (alignment, gene expression quantification and quality control) were consolidated in a final html report generated by the MAPR-seq pipeline. Principal component analysis (PCA) and hierarchical clustering were performed for quality control, exploratory data analysis, and to identify sample outliers. Using the raw gene counts generated from the MAP-RSeq pipeline, genes with significant differential expression between different samples were assessed using edgeR.^41^ Genes with counts per million (CPM) ˂ 0.5 were filtered out prior to differentially expressed genes (DEGs) analysis. Three comparisons were performed including i) Late passage vs. early passage chondrocytes from males with OA; ii) Late passage vs early passage chondrocytes from females with OA and iii) Late passage vs. early passage chondrocytes from healthy males. DEGs were visually represented using a volcano plot and heatmap generated with the R package ggplot2. DEGs were reported along with their magnitude of change (log2 fold change ≥ 1 or ≤ -1) and level of significance (False Discovery Rate, FDR <0.05). Pathway analysis was conducted with R package enrichR^42–44^ or Metascape^45^ (https://metascape.org) using human pathway settings.

### Quantitative Real Time Polymerase Chain Reaction (qRT-PCR)

Total RNA was isolated from samples snap-frozen in liquid nitrogen. First, snap frozen samples were homogenized in an Eppendorf tube using a 3-mm Tungsten carbide beads and TissueLyser II system (Qiagen; 4 min at 30 Hz). RNA isolation steps were followed as mentioned above in this section. 1 µg of cDNA was synthesized using a high-capacity cDNA reverse transcription kit (Applied Biosystems, Waltham, MA, USA). PCR experiments were performed using 20 ng of cDNA per reaction, qRT-PCR Brilliant III ultra-fast master mix kit (Agilent) and TaqMan gene expression probes (Thermo Fisher Scientific) as follows: IL-1α (Hs00174092_m1), IL-1β (Hs01555410_m1), IL-6 (Hs00174131_m1), p16^INK4A^ (Hs00923894_m1), p21^CIP1^ (Hs00355782_m1), p53 (Hs01034249_m1). The PCR assay was conducted using the AriaMx qRT-PCR detection system (Agilent). Gene expression values were normalized to the GAPDH housekeeping gene to calculate the Δ^Ct^ value.

### Enzyme Linked Immunosorbent Assay (ELISA)

IL-1α or IL-1β concentrations in conditioned media were determined using the commercial DuoSet ELISA Kit (R&D Systems, Minneapolis, MN, USA; IL-1α Cat. No.: DY200; IL-1β Cat. No.: DY201).

### Statistical Analysis

The statistical analyses for qRT-PCR and ELISA were performed utilizing GraphPad Prism 9.2.0 (GraphPad, Boston, MA, USA). The experimental design involved 3 biological replicates (n = 3). Data distribution was assessed using either the Kolmogorov–Smirnov test or Shapiro–Wilk tests. A two tailed unpaired t-test or two-way ANOVA was employed for normally distributed data, while the Mann–Whitney U-test was utilized for non-normally distributed data. The results were presented as mean ± standard deviation (SD). Statistical significance was set at a p-value <0.05.

## Results

### Replicative senescence of chondrocytes

Chondrocytes from all three subjects reached their Hayflick limit after 36 doublings (18 passages). Early passage chondrocytes from osteoarthritic donors displayed various cell morphologies, including stellate, elongate and fibroblastic (Figure 2A, C). Late-passage chondrocytes exhibited additional morphological phenotypes, including the large cells shown in Figure 2B, D. Early passage cultures of chondrocytes from the healthy young donor were smaller and comprised a mixture of polygonal and rounded cells (Figure 2E). Terminal passage cells were much larger and displayed a mixture of stellate and flattened phenotypes (Figure 2F).

**Figure 2.**
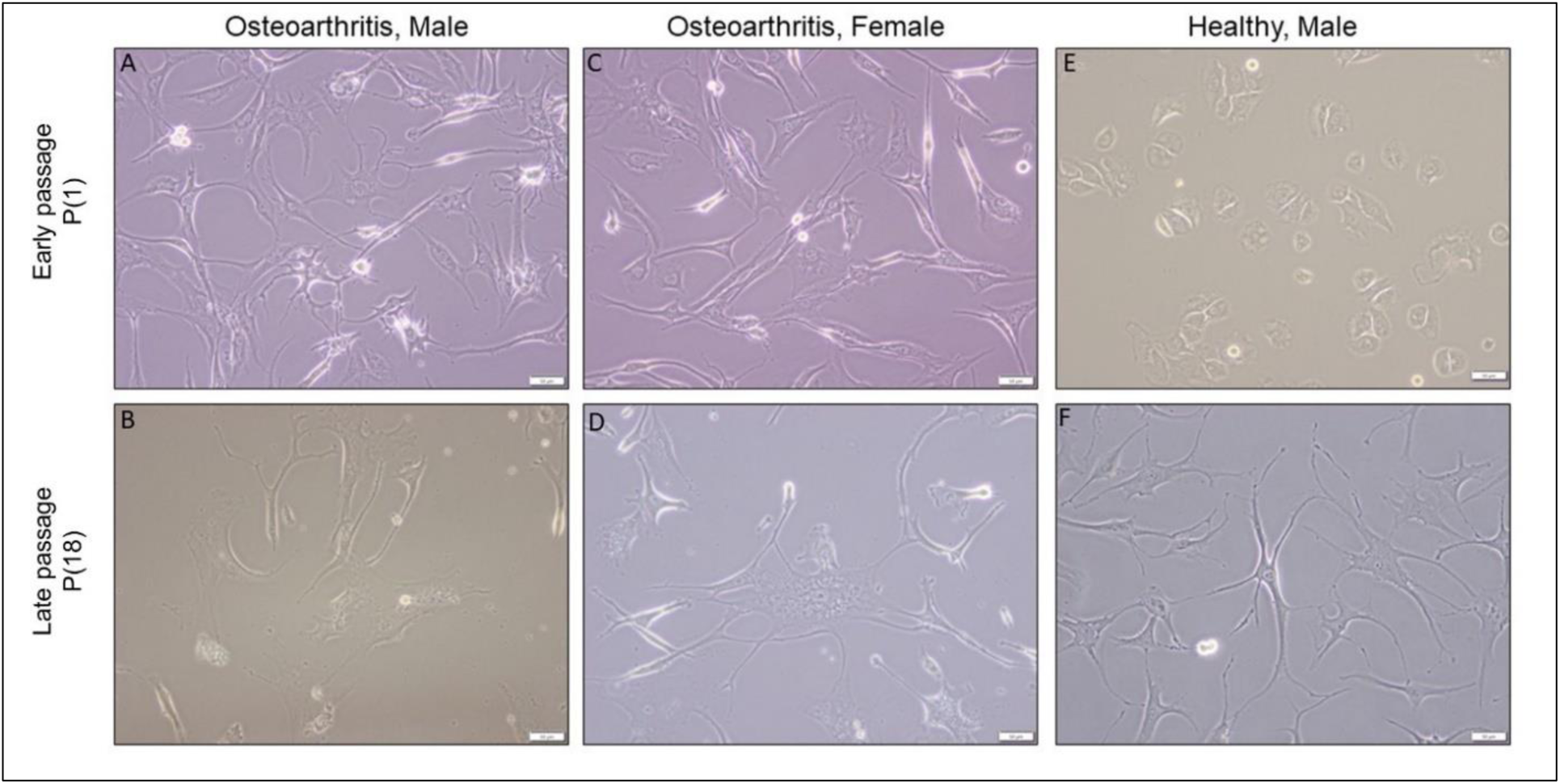
Cell morphology analysis of early and late passage chondrocytes **(A, B)** osteoarthritis, male; **(C, D)** osteoarthritis, female; **(E, F)** healthy, male. The first horizontal row shows early passage chondrocytes, while the second horizontal row displays late passage chondrocytes. Scale bars: 50 µm with 20x obj.

When early and late passage chondrocytes were compared in their ability to form cartilaginous nodules in pellet culture, there were clear differences (Figure 3A). Pellets formed from early passage chondrocytes were larger and displayed the characteristic white, shiny appearance of non-osteoarthritic cartilage. Those formed from late passage cells were smaller with a slight yellow discoloration (Figure 3A). The ECM of pellets formed from late passage chondrocytes lacked the metachromatically staining of pellets formed by early passage cells (Figure 3B).

**Figure 3.**
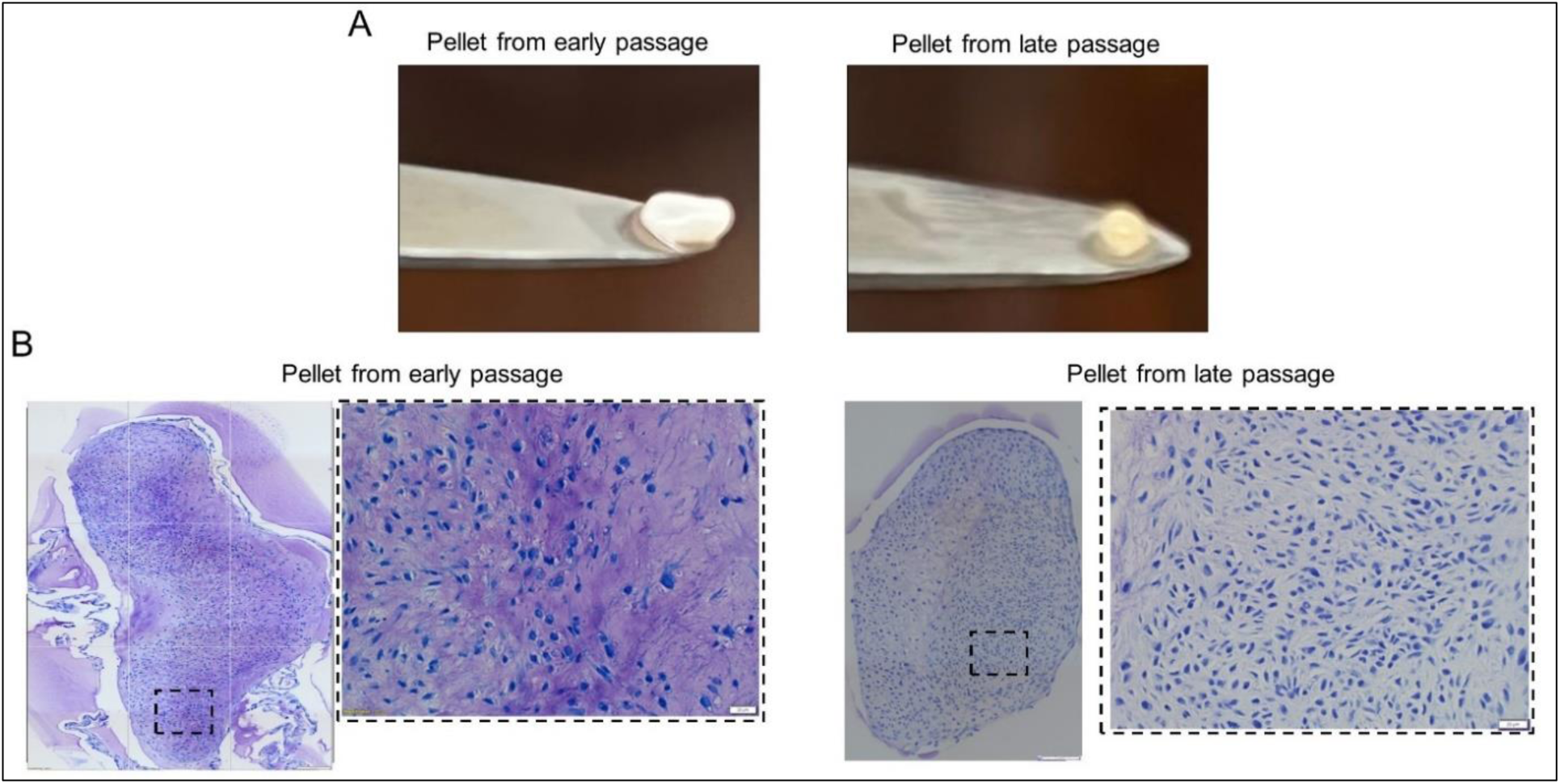
Effects of replicative senescence on the ability of cells to undergo chondrogenesis in 3D chondrocyte pellet culture **(A)** morphology and **(B)** Toluidine blue staining. Scale bar 200µm with 4x objective and 50 µm with 20x objective.

### DEGs between early and late passage chondrocytes

Transcriptomic profiling was performed on early and late passage cultures derived from human articular cartilage samples of all three donors. The primary objective was to examine gene expression changes associated with aging; secondary objectives were to determine the effects of sex and OA, although with small n results are preliminary. PCA of all genes showed clear separation between the late passage and early passage chondrocytes, indicating substantial differences in gene expression (Figure 4). DEGs in early and late passage cells were then analyzed for each of the three chondrocyte cultures.

**Figure 4.**
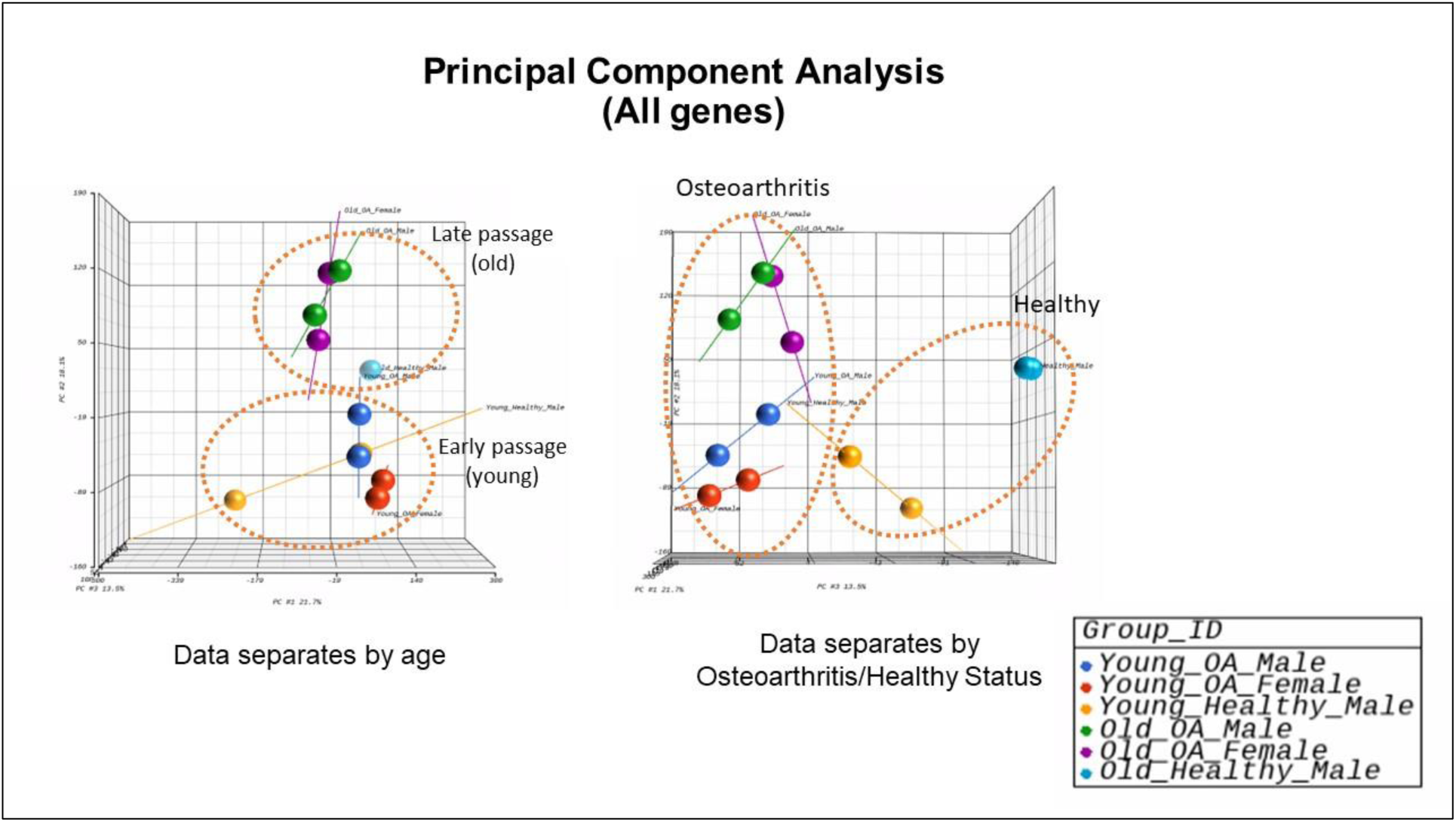
Principal component analysis (PCA) showing sequencing variation of all genes sorted by age (observation from front space) and osteoarthritis/healthy (observation from side space).

#### Sample 1 – Early and late passage chondrocytes from male with OA

Distinct clusters were observed when comparing the transcriptomes of early and late passage chondrocytes obtained from a male with OA (Figure 5A). The heat map clearly demonstrated a notable distinction in the gene expression patterns between early and late passage chondrocytes (Figure 5B). A volcano plot revealed the most significantly altered genes, with ESM1 exhibiting the greatest upregulation and FGD5 showing the greatest downregulation (Figure 5C). The analysis identified 2,309 DEGs, with 983 genes upregulated and 1,326 downregulated in terminal passage chondrocytes (Figure 5D). Pathway analysis revealed downregulation of ECM organization, collagen fibril organization, glycosaminoglycan catabolic process, skeletal and nervous system development, and upregulation of focal adhesion, axon guidance, and DNA replication genes in late passage cells (Figure 5E).

**Figure 5.**
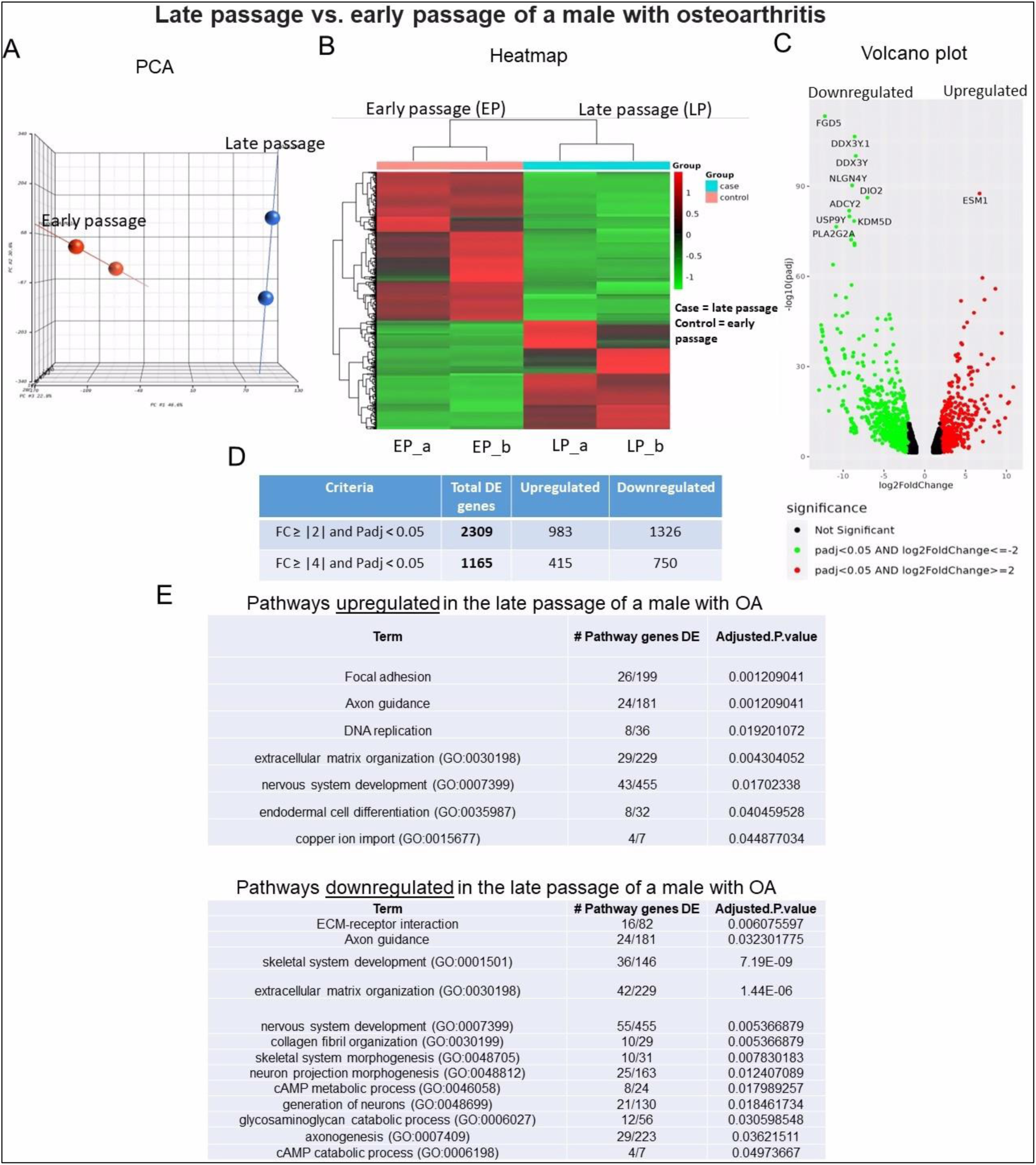
Bulk RNA sequencing analysis of late passage vs early passage chondrocytes from a male with osteoarthritis (sample-1): **A)** Principal component analysis (PCA) indicating differential expression analysis between late and early passage cells; **(B)** Heatmap depicting upregulated (shown as red) and downregulated (shown as green) genes in the late passage vs early passage of chondrocytes. (_a, _b) are duplicates from the same sample; **(C)** Volcano plot highlighting differential gene expression between late passage vs early passage of chondrocytes within samples. Red dots represent significantly differentially expressed genes (DEGs) that have an absolute fold change (FC) of ≥2, green dots represent genes that have an absolute FC of ≤-2, and black dots represent genes not differentially expressed. The FC presented here is the gene expression of late passages relative to early passages; **(D)** Tables showing up/down regulated genes; **(E)** Basic pathway analysis revealing alterations in key pathways linked to aging in late passage showing up/down regulated pathways.

#### Sample 2 – Early and late passage chondrocytes from female with OA

Hierarchical clustering revealed similar distinct transcriptomic signatures to those observed in sample 1 (Figure 6A). This sample displayed a comparable heatmap pattern to that of sample 1 (Figure 6B). The largest gene expression changes in sample 2 were the upregulation of PLAT and the downregulation of SLC7A2 (Figure 6C). A greater number of genes exhibited regulation in sample 2 compared to sample 1 (Figure 6D). Specifically, a total of 3,281 DEGs were identified in the comparison between early and late passage chondrocytes. Among these, 1,577 genes showed upregulation, while 1,704 displayed downregulation. Similar to sample 1, the aging process in this context was associated with the downregulation of genes related to ECM organization and collagen, while genes associated with the cell cycle, DNA replication, and neuron development exhibited upregulation (Figure 6E).

**Figure 6.**
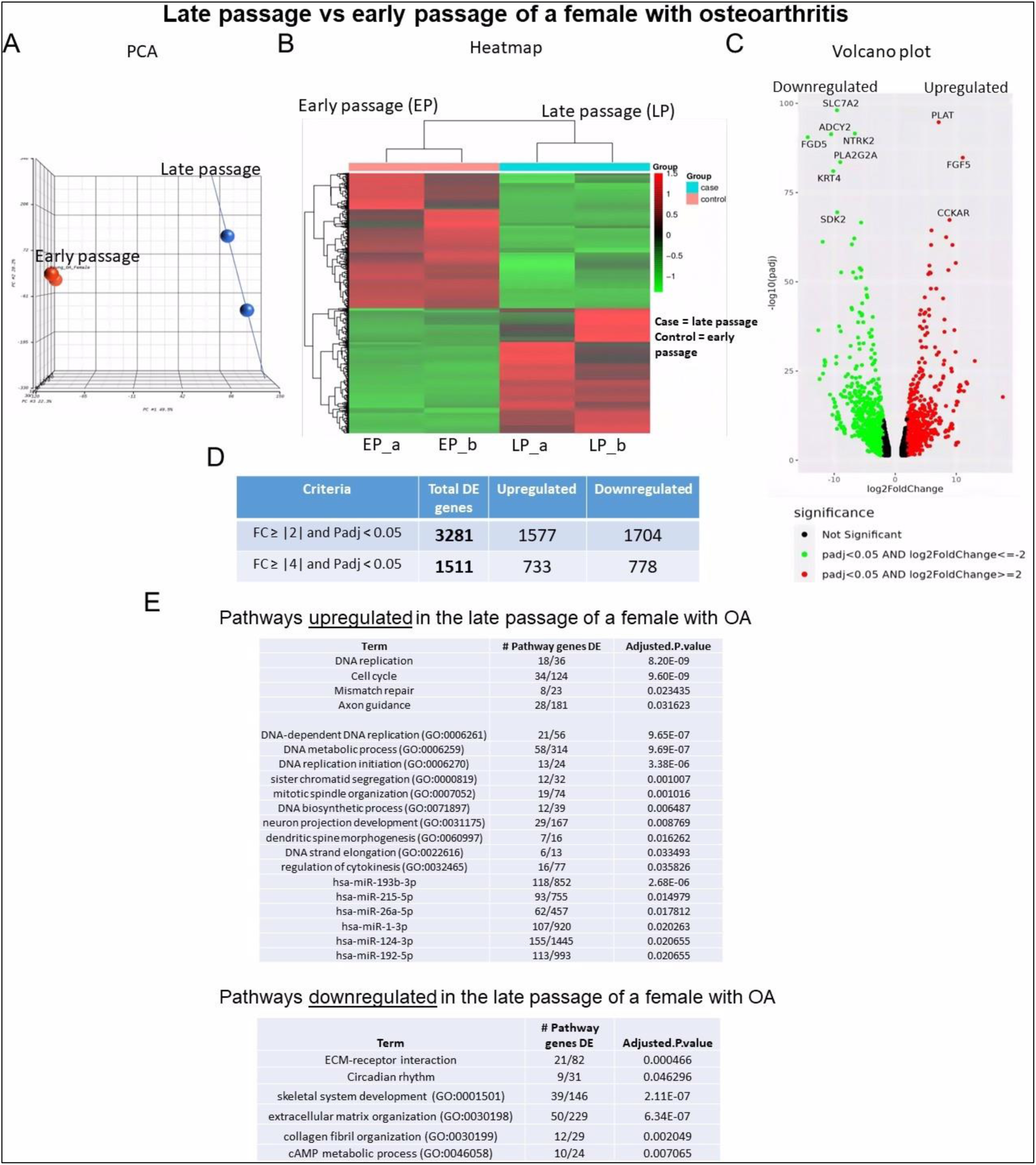
Bulk RNA sequencing analysis of late passage vs early passage chondrocytes from a female with osteoarthritis (sample-2): **A)** Principal component analysis (PCA) indicating differential expression analysis between late and early passage cells; **(B)** Heatmap depicting upregulated (shown as red) and downregulated (shown as green) genes in the late passage vs early passage of chondrocytes. (_a, _b) are duplicates from the same sample; **(C)** Volcano plot highlighting differential gene expression between late passage vs early passage of chondrocytes within samples. Red dots represent significantly differentially expressed genes (DEGs) that have an absolute fold change (FC) of ≥2, green dots represent genes that have an absolute FC of ≤-2, and black dots represent genes not differentially expressed. The FC presented here is the gene expression of late passages relative to early passages; **(D)** Tables showing up/down regulated genes; **(E)** Basic pathway analysis revealing alterations in key pathways linked to aging in late passage showing up/down regulated pathways.

#### Sample 3 - Early vs late passage chondrocytes from a healthy male

Hierarchical clustering in this comparison revealed a less distinct pattern, particularly when contrasted with samples 1 and 2 (Figure 7A). Additionally, the heatmap displayed an opposing pattern in this comparison, differing from the observed patterns in sample 1 and 2 (Figure 7B). The volcano plot highlighted SLC7A2 as the most downregulated gene, while CLCA2 appeared as the most upregulated gene (Figure 7C). In contrast to the comparison between samples 1 and 2, fewer DEGs (1,298) were identified in this sample (Figure 7D), suggesting more modest changes associated with senescence without OA. Notably, cell cycle and DNA replication genes exhibited downregulation, while prostaglandin genes were upregulated in old males (Figure 7E).

**Figure 7.**
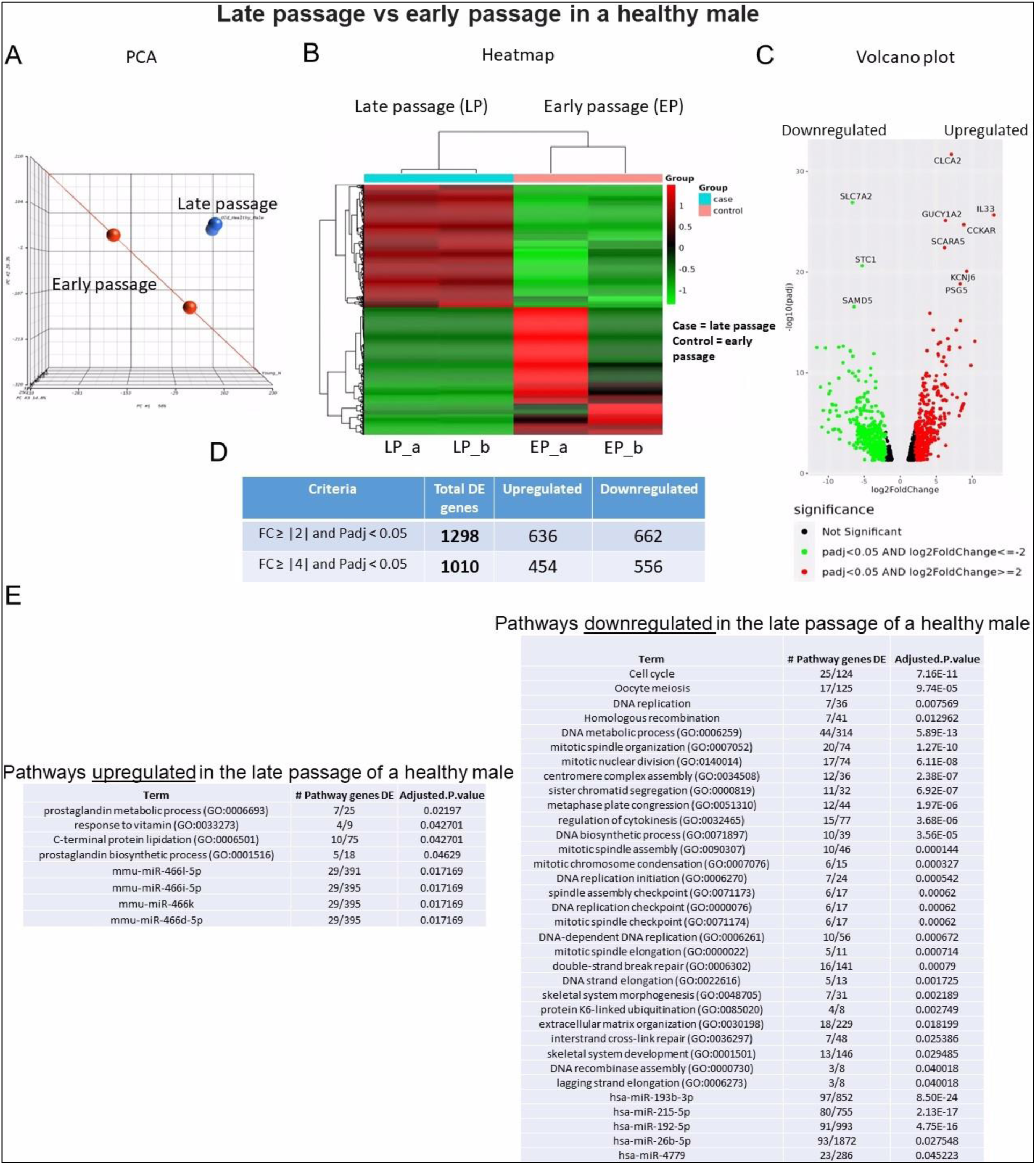
Bulk RNA sequencing analysis of late passage vs early passage chondrocytes from a healthy male (sample-3): **A)** Principal component analysis (PCA) indicating differential expression analysis between late and early passage cells; **(B)** Heatmap depicting upregulated (shown as red) and downregulated (shown as green) genes in the late passage vs early passage of chondrocytes. (_a, _b) are duplicates from the same sample; **(C)** Volcano plot highlighting differential gene expression between late passage vs early passage of chondrocytes within samples. Red dots represent significantly differentially expressed genes (DEGs) that have an absolute fold change (FC) of ≥2, green dots represent genes that have an absolute FC of ≤-2, and black dots represent genes not differentially expressed. The FC presented here is the gene expression of late passages relative to early passages; **(D)** Tables showing up/down regulated genes; **(E)** Basic pathway analysis revealing alterations in key pathways linked to aging in late passage showing up/down regulated pathways.

#### Analysis of ECM components, enzymes that degrade ECM and SASP factors

A focused analysis was undertaken on components of the ECM, the enzymes that degrade them and SASP factors within each sample (Figure 8). All samples showed marked downregulation of Col2A1 and ACAN in late passage chondrocytes (Figure 8A). Conversely, MMP19 demonstrated a notable upregulation in the late passage chondrocytes of all samples. Expression of ADAMTS4 and ADAMTS8 showed significant upregulation in late passage chondrocytes obtained from OA samples, but not in those obtained from healthy cartilage.

**Figure 8.**
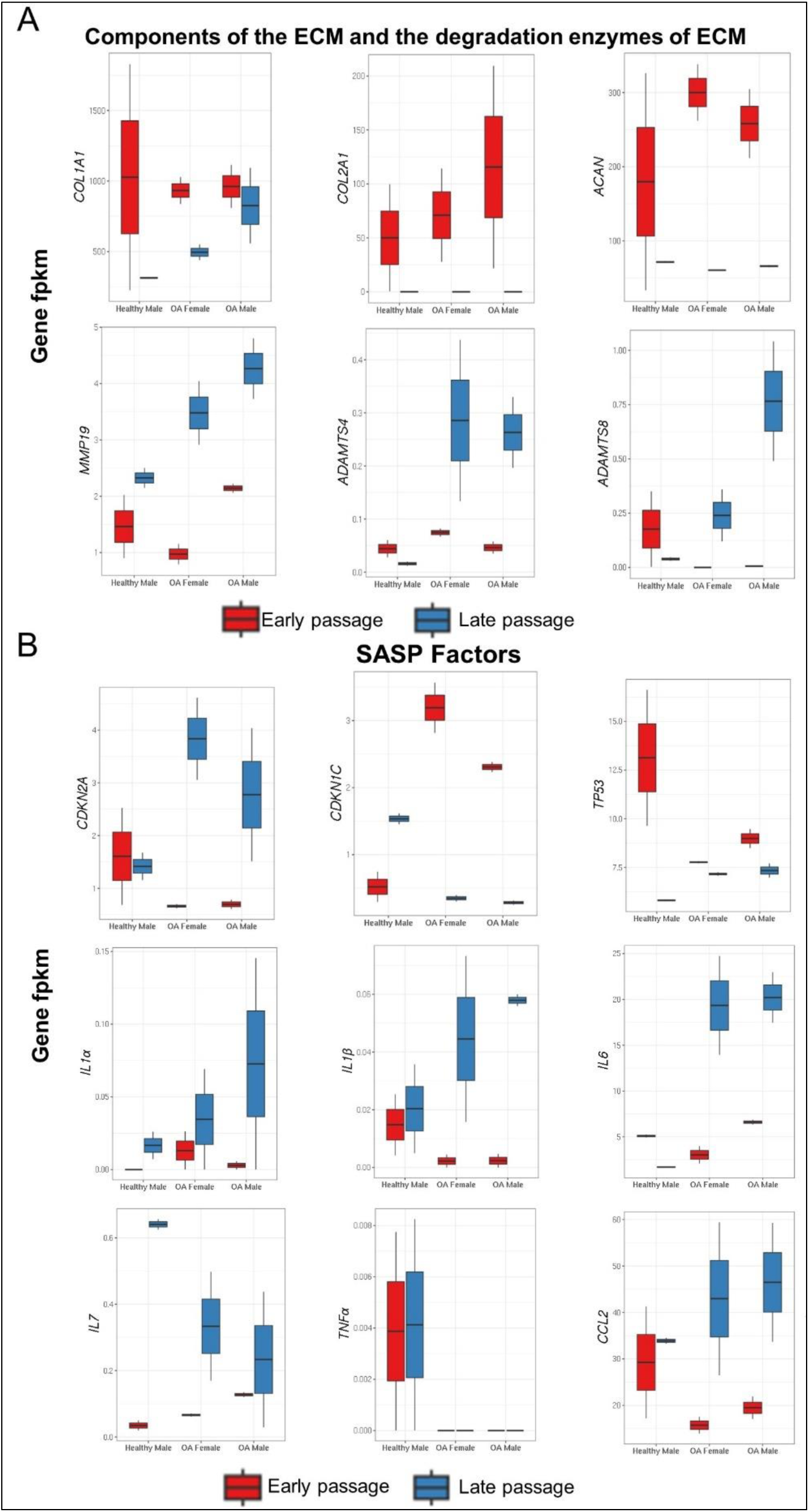
DEGs encompassing **(A)** components of the extracellular matrix (ECM), enzymes and that degrade the ECM, and **(B)** genes associated with the senescence-associated secretory phenotype (SASP). (Gene fpkm: Fragments Per Kilobase per Million mapped reads).

Among transcripts associated with the SASP, CDKN2A (p16^INK4A^); CKN1C: (p21^CIP1^); TP53: (p53), only p16^INK4A^ demonstrated a significant upregulation in late passage of chondrocytes within the OA samples compared to healthy samples (Figure 8B). IL-1α and IL-7 expression increased in all late passage samples, whereas IL-1β and IL-6 transcripts only increased in late passage cells derived from OA samples. There was no change in TNFα expression with passage in any of the samples. CCL2 was upregulated in the late passage of chondrocytes derived from patients with OA, but not those derived from a healthy subject.

### Pathway analysis comparing male and female with OA

Pathway analysis was conducted to compare the DEGs profiles between chondrocytes derived from the male and female samples with OA, utilizing data from samples 1 and 2. This analysis revealed 611 genes unique to males and 1,583 genes unique to females, with 1,698 genes common between the sexes (Figure 9A). The top 20 pathways <u>common</u> between males and females with OA included ECM organization, collagen formation, musculoskeletal development including bone and muscle, and nervous system development (Figure 9B). The top 20 pathways unique to <u>males</u> with OA encompassed ECM organization, mitotic cell cycle, and regulation of neuron differentiation (Figure 9C). The top 20 pathways unique to <u>females</u> with OA featured in regulation of cell cycle process, DNA metabolic process, and PID-PLK1 pathway (Figure 9D).

**Figure 9.**
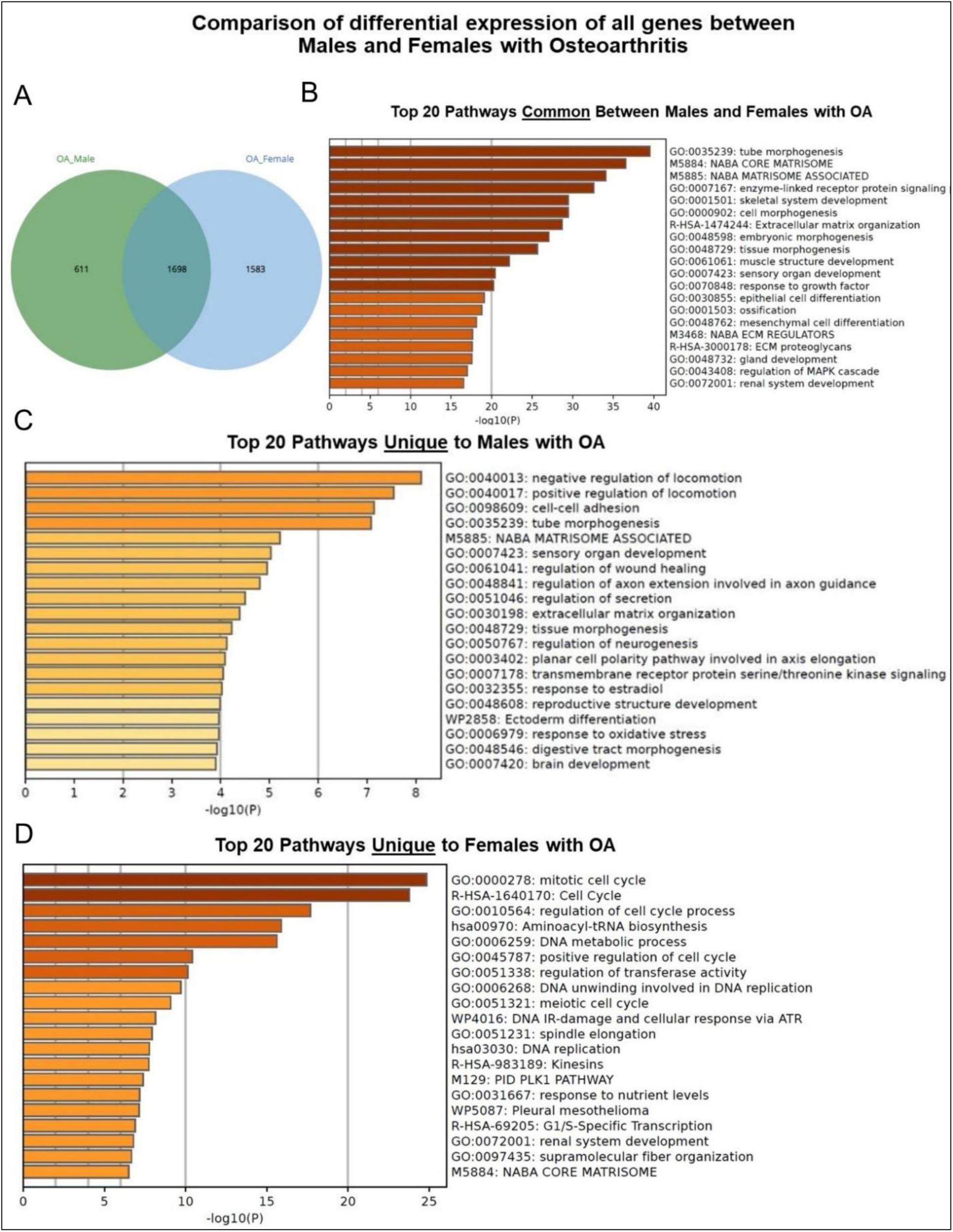
Basic pathway analysis of differentially expressed genes (DEGs) of males and females with osteoarthritis **(A)** Venn diagram illustrating shared and distinct genes in males and females; **(B)** Top 20 Pathways <u>Common</u> Between Males and Females with OA; **(C)** Top 20 Pathways <u>Unique to Males</u> with OA; **(D)** Top 20 Pathways <u>Unique to Females</u> with OA.

### Quantitative RT-PCR and ELISA

Certain findings from bulk RNA sequencing were validated by qRT-PCR (Figure 10). Consistent with the RNA sequencing data, significant upregulation of CDKN2A (p16^INK4A^), IL-1α, IL-1β, and IL-6 was observed in late passage chondrocytes (Figure 10A). Increased levels of IL-1α and IL- 1β protein were confirmed by ELISA (Figure 10B).

**Figure 10.**
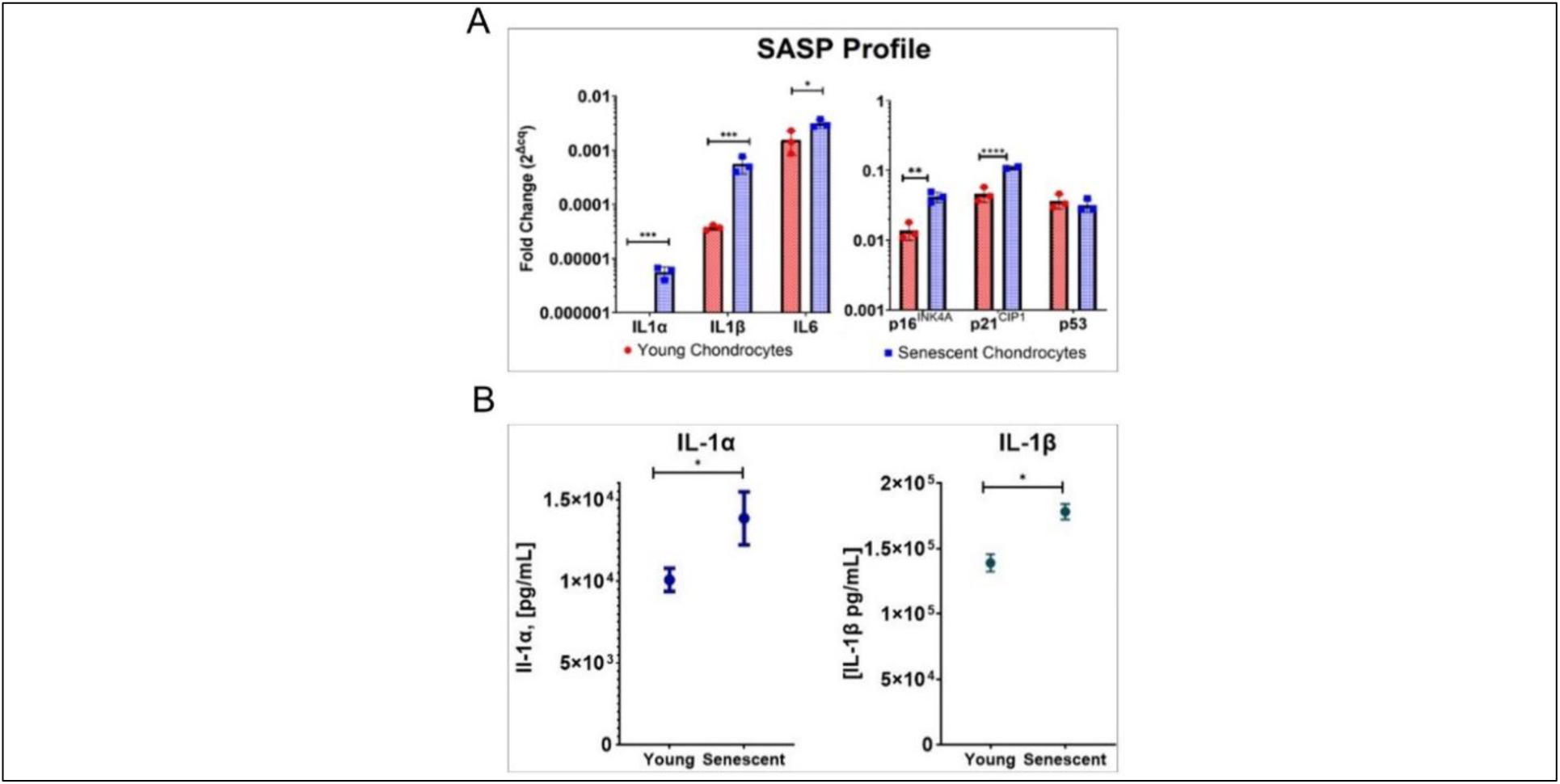
qRT-PCR and ELISA data. **(A)** qRT-PCR analysis of IL-1α, IL-1β, IL-6, and p16^INK4A^, p21^CIP1^, and p53; **(B)** Protein concentrations of IL-1α and IL-1β. (*p<0.05; **p<0.01; ***p<0.001; ****p<0.0001).

## Discussion

In this study, we demonstrated that serial passaging of chondrocytes to replicative senescence induced characteristic morphological changes and reduced chondrogenic potential. Chondrocytes reached replicative senescence at passage 18 after 36 population doublings. It is remarkable that chondrocytes derived from old (72 years, 80 years) and young (26 years) donors had the same Hayflick limit, as diploid cells from young individuals typically have a greater mitotic potential than those from old individuals. It is possible that the quiescent state of articular chondrocytes *in vivo* preserves their mitotic potential. Nevertheless, when plated into monolayer culture, chondrocytes from the two old donors already had the morphology of late passage cells. Because this is a small pilot study involving only 3 donors it is not possible to draw strong conclusions, but these observations are nevertheless intriguing. We also showed that late passage chondrocytes have a reduced capacity to form cartilaginous tissues, as evidenced by smaller pellet sizes and decreased proteoglycan content. This indicates replicative senescence impairs the chondrogenic potential of cultured chondrocytes.

Bulk RNA sequencing, conducted on chondrocytes from each donor, revealed clear separation between early and late passage cells in PCA, highlighting substantial differences in gene expression. The heatmap representing the chondrocytes obtained from joints with OA (Sample 1 & 2) exhibited a pattern from that of chondrocytes harvested from a healthy joint (Sample 3). Volcano plots revealed that FGD5 was the most downregulated gene in late passage chondrocytes derived from male, OA cartilage (Sample 1). While the precise involvement of FGD5 in OA development is not entirely elucidated, Yang et al. ^46^ demonstrated that FGD5-AS1 regulates OA progression by miR-302d-3p/TGFBR2 axis, protecting chondrocytes from inflammation-induced injury and reducing the ECM degradation. Consistent with this, SLC7A2 was strongly downregulated in late passage chondrocytes from both female OA cartilage and healthy male cartilage (Sample 2 & Sample 3). Ni et al. ^47^ showed the protective role of circular SLC7A2 against OA through inhibition of the miR-4498/TIMP3 axis. CLCA2 was the most upregulated genes in late passage chondrocytes derived from healthy male cartilage (Sample 3). Our findings agree with those of Tanikawa et al. ^48^ who found that CLCA2 is significantly induced during the replicative senescence of mouse embryonic fibroblasts, particularly as a p53-inducible mediator of senescence.

Expression of Col2A1 and ACAN declined in late passage cells, while MMP19, ADAMT4 and ADAMT8 were upregulated, consistent with reduced integrity of the ECM with senescence. The pro-inflammatory cytokines IL-1α and IL-7 were upregulated in late passage chondrocytes from both OA and healthy samples. However, the transcription factor p16^INK4A^ and the pro-inflammatory cytokines IL-1β and IL-6 were upregulated specifically in late passage chondrocytes from OA samples compared to healthy samples, along with the chemokine CCL2. In contrast, TNFα was undetectable in nearly all samples and did not display any change in expression with passage.

Confirmation of RNA sequencing data by qRT-PCR of certain transcripts further confirmed the observed gene expression changes. The upregulation of p16^INK4A^, IL-1α, IL-1β, and IL-6, along with elevated protein concentrations of IL-1α and IL-1β, further supported the association between senescence and OA. While p16^INK4a^ is an important cell cycle inhibitor, it is not needed for SASP production. Diekman et al. ^49^ showed that knocking out p16^INK4a^ in chondrocytes of adult mice did not reduce SASP expression or change the rate of OA. These findings show that although p16^INK4a^ expression marks dysfunctional chondrocytes, the effects of chondrocyte senescence on OA progression are more likely mediated by SASP production rather than loss of chondrocyte proliferation. IL-1β is one of the most potent stimulators of cartilage breakdown. It inhibits synthesis of type II collagen and proteoglycans and promotes the synthesis of MMPs, ADAMTS^50^ and IL-6;^51^ it is interesting that expression of TNFα was not detected in most samples. Our data also showed IL-7 as another proinflammatory cytokine whose expression increases in late passage cultures. Consistent with this, Long et al. ^24^ demonstrated that human articular chondrocytes in monolayer culture derived from older donors secrete elevated levels of IL-7 compared to their younger counterparts. They further showed that chondrocytes from OA patients release higher amounts of IL-7 than age-matched individuals with normal joint health.

To identify sex specific DEGs in OA, basic pathway analysis was performed. The top 20 pathways common to both males and females with OA were generally related to ECM organization, collagen formation, musculoskeletal development including bone and muscle, and nervous system development. However, females (1583 genes) showed distinct pathways compared to males (611 genes), including regulation of cell cycle processes, DNA metabolism, and the PLK1 signaling pathway. PLK1 is a key regulator of mitotic cell division and has been shown to be overexpressed in several human cancers, including breast cancer. ^52^ Driscoll et al. ^53^ showed that inhibiting PLK1 causes post-mitotic DNA damage and senescence in tumor cell lines. Kim et al. ^54^ found that decreasing PLK1 levels stimulates senescence through a p53-dependent pathway. Additionally, Wierer et al. ^55^ showed that PLK1 mediates estrogen receptor-regulated gene in human breast cancer cells. Taken together, PLK1 is an important regulator of cell cycle progression, senescence, and estrogen receptor signaling, suggesting PLK1 may play a key role in the sex differences seen in OA pathology, especially through its effects on responses to estrogen.

Small sample size is the main limitation of this pilot study. Additional studies with more subjects from both sexes are important to fully characterize chondrocyte senescence and elucidate subtle sex differences in relation to its pathophysiologic role in OA. The limited availability of normal cartilage specimens is a major obstacle to achieving this in a fast fashion. Another consideration is collecting samples from individuals with idiopathic OA, where chondrocyte senescence may be the key underlying pathophysiologic process. Overall, while preliminary, these data provide insights into transcriptomic changes underlying chondrocyte aging and OA development. Further research with larger sample sizes representing defined cohorts of subjects will allow more robust characterization of the complex molecular interplay between aging, chondrocyte senescence, sex and OA.

## Conclusion

In this small (n=3) pilot study, clear distinctions in DEGs were observed between early and late passage chondrocytes, as well as between males and females and those with or without OA. These findings indicate possible important age- and sex-related differences in gene expression associated with OA. The upregulation of IL-1α, IL-1β, IL-6, and IL-7 and p16^INK4a^, in late passage chondrocytes from OA samples highlights the link between cellular senescence and inflammation. This was associated with significant downregulation of key ECM genes Col2A1 and ACAN, along with upregulation of matrix-degrading enzymes in late passage chondrocytes derived from joints with OA, consistent with a pivotal role of chondrocyte senescence in OA pathogenesis. Additionally, the sample derived from a female patient exhibited DEG patterns that differed from those of samples derived from males, although more samples are needed to confirm a sex-related difference in DEG. Further research with larger cohort sizes is necessary to fully characterize the complex interplay between chondrocyte senescence, aging, sex and OA development.

## Data availability

All computational analysis pipelines can be found on NCBI Gene Expression Omnibus (GEO) (https://www.ncbi.nlm.nih.gov/geo/) under accession numbers GSE246425.

## Author Contributions

AAZ; study design, data analysis and interpretation, writing original draft. GH; data acquisition. CN: data acquisition, review & editing. CLDP: histology, review & editing. DS: resources, review & editing. MA: resources, review & editing. CHE: Conceptualization, investigation, review & editing, supervision. All authors have read and agreed to the published version of the manuscript.

## Acknowledgements

RNA sequencing was conducted in collaboration with the Mayo Clinic Genome Analysis Core. Bioinformatics support was provided by the Department of Quantitative Health Sciences. Graphical abstract was created with BioRender.com.

